# PlotsOfDifferences – a web app for the quantitative comparison of unpaired data

**DOI:** 10.1101/578575

**Authors:** Joachim Goedhart

## Abstract

The quantitative comparison of data acquired under different conditions is an important aspect of experimental science. The most widely used statistic for quantitative comparisons is the p-value. However, p-values suffer from several shortcomings. The most prominent shortcoming that is relevant for quantitative comparisons is that p-values fail to convey the magnitude of differences. The differences between conditions are best quantified by the determination of effect size. To democratize the calculation of effect size, we have developed a web-based tool. The tool uses bootstrapping to resample mean or median values for each of the conditions and these values are used to calculate the effect size and their compatibility interval. The web tool generates a graphical output, showing the bootstrap distribution of the difference next to the actual data for optimal interpretation. A tabular output with statistics and effect sizes is also generated and the table can be supplemented with p-values that are calculated with a randomization test. The app that we report here is dubbed PlotsOfDifferences and is available at: https://huygens.science.uva.nl/PlotsOfDifferences

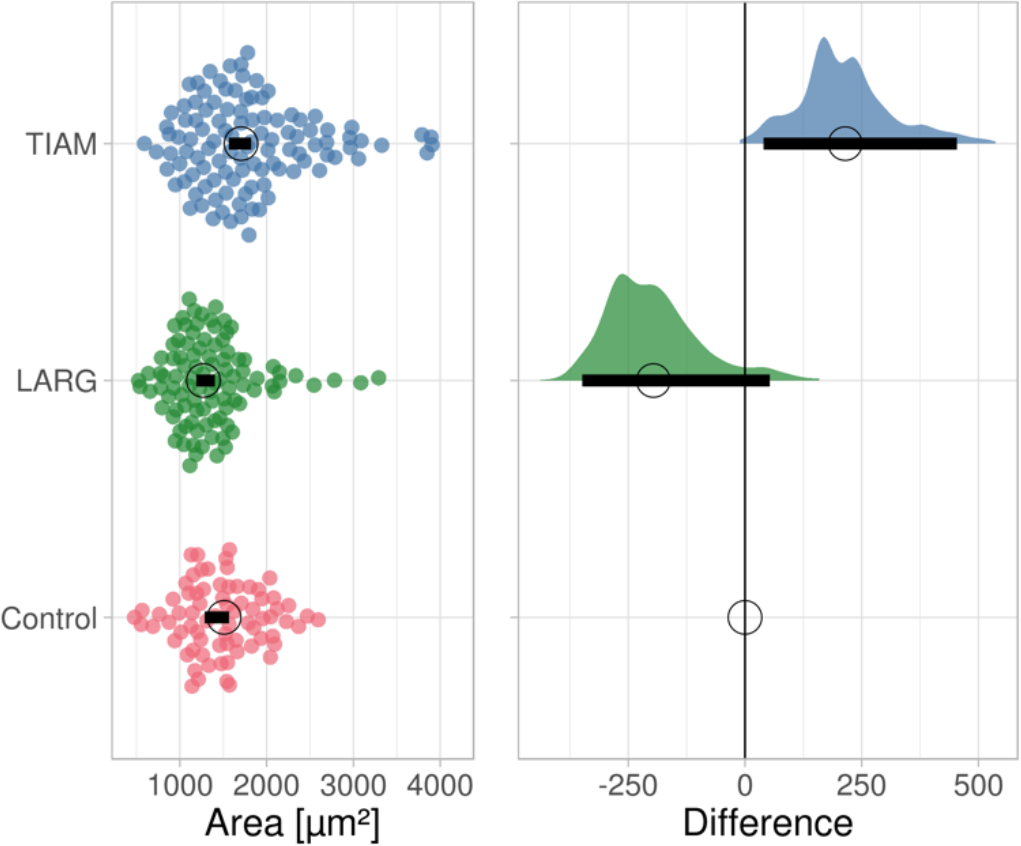

## Introduction

Performing measurements under different conditions is a classic strategy in scientific experimentation. Usually one of the conditions is a reference or control condition and to the other conditions systematic perturbations are applied. To examine the effect of the perturbation, it is necessary to compare the data obtained under different conditions. A graphical representation of the data provides a straightforward and powerful way for evaluating differences and similarities (Drummond and Vowler, 2011). Statistics can be used to facilitate the comparison. A direct, but qualitative comparison of the different categories/conditions can be done by ‘visual inference’ if 95% confidence intervals are supplied (Cumming and Finch, 2005; Cumming et al., 2007; Gardner and Altman, 1986).

Yet, in many cases a quantitative or objective comparison is desired. The standard practice is to subject the data to a statistical test that results in a p-value. This method is so entrenched in the routine of data analysis and presentation that it is often applied when it is superfluous (Goedhart, 2018). Another issue is that p-values are poorly understood due to its non-intuitive definition, resulting in misinterpretation and misuse (Goodman, 2008). Finally, the p-value does not reflect the magnitude of a difference between conditions, whereas the difference is usually the measure of interest (Drummond and Tom, 2011).

The difference between conditions is reflected by the ‘effect size’. Many different ways of calculating the effect size have been suggested and they are roughly classified in relative and absolute effect sizes. The reporting of effect sizes is rare in the biomedical sciences and examples are difficult to find, despite a clear advantage of effect size over significance testing (Claridge-Chang and Assam, 2016; Cumming, 2014; Gardner and Altman, 1986). In cases where effect size is reported (Mohammad et al., 2017), custom written scripts were used to calculate and display the effect size in graphs. Thus, a lack of user-friendly tools for the calculation and presentation of effect sizes hinders the wide adoption of effect sizes for the quantitative comparison of data. Recently, easy-to-use tools for calculation of relative effect size (Goedhart, 2016) and absolute effect sizes have been reported (Ho et al., 2018) and these tools are important for wide adoption of effect sizes. We have previously reported a web tool, PlotsOfData, for visualizing the actual data with summary statistics (Postma and Goedhart, 2018). Here, we report an extension of that web tool that adds the calculation and presentation of differences and their compatibility interval for the quantitative comparison of conditions. For educational reasons, a p-value can be provided as well. The p-value calculated by the web tool are based on the randomization method which is intuitive and makes no assumption about the underlying data distribution (Nuzzo, 2017;Hooton, 1991).

## Availability & code

The webtool is available at: https://huygens.science.uva.nl/PlotsOfDifferences

The app uses the shiny package and was written in R using R (https://www.r-project.org) and Rstudio (https://www.rstudio.com). It uses several freely available packages (shiny, ggplot2, dplyr, tidyr, readr, magrittr, ggbeeswarm, readxl, DT, gridExtra). The current version 1.0.0 is available at zenodo: https://doi.org/10.5281/zenodo.2573907. The up-to-date source code is available at Github together with information on how to install and run the app locally:https://github.com/JoachimGoedhart/PlotsOfDifferences

## Data input and structure

The data can be supplied in different ways, similar to the input for PlotsOfData (Postma and Goedhart, 2018). Both wide and tidy (Wickham, 2014) data structures are accepted. The wide format is used as a default, but it can be changed to tidy by using an alternative hyperlink: https://huygens.science.uva.nl/PlotsOfDifferences/?data=4;T

## Plotting the data

A detailed description of the options for plotting the data is reported in the paper on PlotsOfData (Postma and Goedhart, 2018).

## Plotting the effect size

A bootstrap method is used to calculate the effect size and the compatibility interval (CI). To this end, each condition is resampled 1000x with replacement to calculate a distribution of mean or median values. A reference condition is selected by the user and the difference between the collection of boostrapped median or mean values is calculated, resulting in a new distribution of differences. This distribution is plotted. The 2.5^th^ and 97.5^th^ percentile of the distribution are used to determine a 95% confidence interval around the difference. We will indicate this interval as the compatibility interval (CI), i.e. the range of values most compatible with the data. The CI is displayed as black line under the distribution.

The effect size plot is either displayed next to the data plot (landscape orientation) or below the data plot (portrait orientation). Two file formats are available for downloading the figure, PDF and PNG. The PNG format is lossless can be readily converted to other bitmap-type formats that are suitable for presentation or incorporation into (multi-panel) figures. The PDF format is vector-based and can be imported into any software package that handles vector-based graphics for further adjustment of the lay-out.

## Randomization test

To calculate a p-value of the different conditions relative to the reference, a randomization test is used. Briefly, the data of a condition and the reference are combined (since the implication of the null-hypothesis is that the data are sampled from the same population). The combined data are resampled without replacement and a new (null-) distribution of the difference between means or medians is obtained. To derive a p-value for a two-tailed test, the absolute difference between means or medians is compared with the absolute differences that compose the null-distribution.

## Table with Statistics

A customizable table with statistics is generated, as described previously (Postma and Goedhart, 2018). In addition, a table with effect sizes and their CI is generated. The tables can be exported in different formats (CSV, XLS and PDF).

**Figure 1:**
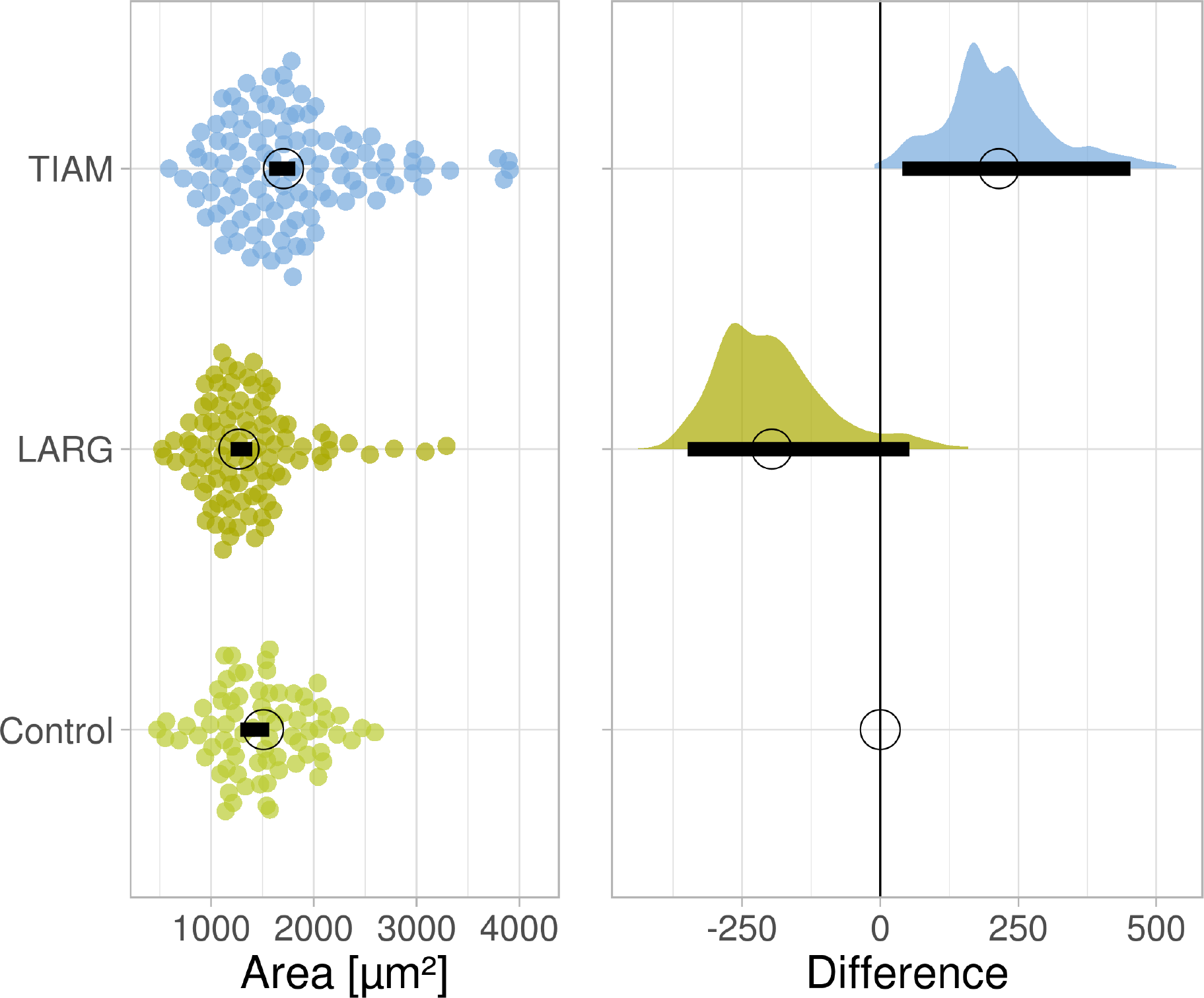
Example graphical output of PlotsOfDifferences for the analysis of difference between median cell area. The difference between median values is determined relative to the ‘Control’ condition and indicate with a circle. The compatibility interval is derived from the bootstrap distribution and indicated with the horizontal bar.

## Application

To illustrate the output of the web-tool, we used data from cell area measurements. The area of cells was measured under a reference, unperturbed condition and two conditions where a Guanine Exchange Factor (GEF) was overexpressed. Plots of the data show that the median values of the perturbed conditions differ from the reference (indicated as ‘Control’). Quantification of the difference between the reference and the other two conditions is shown in the right panel of the figure. The table with values is also presented. The median area of the ‘TIAM’ condition is larger by 215 μm^2^ and the area of the ‘LARG’ condition is smaller by 196 μm^2^. The distribution of differences from the bootstrap procedure is shown and the compatibility interval (CI) that is based on the distribution is indicated with the horizontal black bar. The CI of the effect of TIAM [40 μm^2^ − 454 μm^2^] is the range of values that is most compatible with the data. The CI of the effect of LARG [−349 μm^2^ − 53 μm^2^] does include zero (suggesting no effect) but any effect size in the compatibility interval is conceivable. Therefore, it cannot be concluded that LARG has an effect on area size and it cannot be excluded as well. The effect of LARG on cell area can only be evaluated after acquiring more data, thereby decreasing the uncertainty of the effect size.

The p-values from the randomization test are listed in the table. The p-value for the difference between medians for the TIAM condition is 0.026 and that for LARG is 0.015. Both indicate that the observed difference is unlikely given the hypothesis that the samples (condition and perturbation) were acquired from the same population. For TIAM the p-value is compatible with the effect size, while the p-value for LARG seems on the low side, given that the CI of the effect size includes 0. This stresses the careful evaluation of the outcome of statistical tests in relation to the actual data that is acquired.

**Figure 2:**
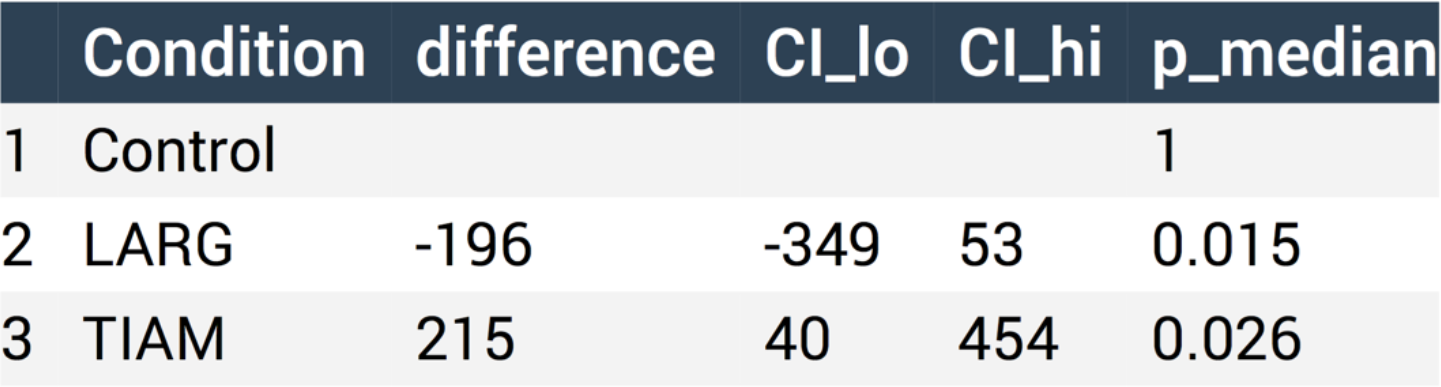
Example tabular output of PlotsOfDifferences for the analysis of difference between median cell area. The difference is determined relative to the ‘Control’ condition.

## Conclusion

A shiny based webtool that uses R without the need for coding skills was generated to democratize quantitative data comparison by calculating absolute differences. The calculated difference is an absolute effect size that is a good alternative (or supplement) for null-hypothesis significance tests and p-values (Halsey et al., 2015; Wasserstein and Lazar, 2016; Claridge-Chang and Assam, 2016; Drummond and Tom, 2011).

A limit of the bootstrap approach is that will not be valid for small sample size (n<10). The premise of bootstrapping is that the sample reflects the population. Obviously, it is difficult to ensure this for low n and the data will better reflect the population for high n. We propose a cut-off at a sample size of 10. The user will receive a warning for n<10 but the webtool will still calculate the effect size and p-value for educational purposes. A feature of the random resampling that is used for calculation of the effect size and the randomization test is that the results will slightly vary between repeated calculations on the same data.

To conclude, we anticipate that the high-quality plots created with PlotsOfDifferences will facilitate quantitative comparisons and improve transparent communication of scientific data which will be beneficial for both researchers and their audience.

## Acknowledgments

The colorblind safe palettes were developed by Paul Tol (https://personal.sron.nl/~pault/). We are grateful to Auke Folkerts (UvA, The Netherlands) for help with the server that runs shiny.

